# Computational prediction shines light on type III secretion origins

**DOI:** 10.1101/036251

**Authors:** Tatyana Goldberg, Burkhard Rost, Yana Bromberg

## Abstract

Type III secretion system is a key bacterial symbiosis and pathogenicity mechanism responsible for a variety of infectious diseases, ranging from food-borne illnesses to the bubonic plague. In many Gram-negative bacteria, the type III secretion system transports effector proteins into host cells, converting resources to bacterial advantage. Here we introduce a computational method that identifies type III effectors by combining homology based inference with *de novo* predictions, reaching up to 3-fold higher performance than existing tools. Our work reveals that signals for recognition and transport of effectors are distributed over the entire protein sequence instead of being confined to the N-terminus, as was previously thought. Our scan of hundreds of prokaryotic genomes identified previously unknown effectors, suggesting that type III secretion may have evolved prior to the archaea/bacteria split. Crucially, our method performs well for short sequence fragments, facilitating evaluation of microbial communities and rapid identification of bacterial pathogenicity – no genome assembly required. pEffect and its data sets are available at http://services.bromberglab.org/peffect.

## Introduction

Six secretion systems have been identified in pathogenic and endosymbiotic Gram-negative bacteria^1–6^. The type III secretion system (T3SS) mediates a wide range of bacterial infections in human, animals, and plants^7^. This system comprises a hollow needle-like structure localized on the surface of bacterial cells that injects specific bacterial proteins, effectors, directly into the cytoplasm of a host cell^3^. During infection, effectors act in concert to convert host resources to their advantage and promote pathogenicity^8^. While the elements of T3SS are conserved among different pathogens, effector proteins are not^7,9,10^.

Although, next generation sequencing techniques yield an ever-growing number of bacterial genome sequences ^11^, experimental verification needed to identify type III effectors remains very expensive and time-consuming. Considering the central role these proteins play in pathogenicity and symbiosis, there is a need for computational tools that predict and prioritize type III effector proteins. To address this need various machine-learning algorithms^12–15^ have been developed to identify type III effectors *in silico*. As input, these methods use similarities in gene GC content and protein amino acid composition, secondary structure, and solvent accessibility to experimentally known effectors. Most often only the protein N-terminus is considered, as it is assumed to be most informative for the translocation of effectors through the type III secretion process ^16^. An independent benchmark revealed that state-of-art-methods predict type III effectors at similar levels of up to 80% accuracy at 80% coverage^17^; thus, there is still room for substantial improvement.

We built pEffect, a method that combines two components -sequence similarity-based inference (PSI-BLAST^18^) and *de novo* prediction using Support Vector Machines (SVM^19^). Our method attains 87±7% accuracy at 95±5% coverage in predicting type III effectors, significantly outperforming each of its components. It also provides a score reflecting the strength of each prediction, allowing users to focus on most relevant results. When tested on sequence fragments similar in length to peptides translated from shotgun sequencing reads, pEffect’s performance was not significantly different. This result suggests that the information required for distinguishing effectors is not confined to any particular part of the amino acid sequence.

We applied pEffect to complete proteomes of over 900 prokaryotic species. pEffect’s high prediction accuracy and ubiquitous applicability raises an interesting question about its predictions of effectors in Gram-positive bacteria and archaea, which are not known to utilize type III secretion. For bacteria, these findings may hint at shared ancestry between flagellar and type III secretion systems^9^. Gene genealogies^20^ and protein network analysis approaches^21^ suggest evolution of both systems from a common ancestor. For archaea common ancestry is less clear. However, predominance in the number of predicted effectors in Gram-negative bacteria, as opposed to the number of predicted effectors in Gram-positive bacteria and archaea suggests repurposing of effector-like proteins independent of organism secretory abilities. In addition to pEffect’s application to evolutionary inference, we find that the time and T3SS completeness–driven results, which follow expectations for correlation with quantities of predicted effectors, are reassuring of our method’s performance.

Our method provides a basis for rapid identification of T3SS-utitlizing bacteria and their exported effector proteins as targets for future therapeutic treatments. The method also proposes interesting directions in which the evolution of bacterial pathogenicity can further be explored. Finally, we suggest using pEffect as a starting point for studies of interactions within microbial communities, detected directly from metagenomic reads and without the need for individual genome assembly.

## Results

### pEffect succeeded linking homology-based and *de novo* predictions

Most functional annotations of new proteins originate from homology-based transfer, *i.e.* on the basis of shared ancestry with experimentally characterized proteins^22^. For type III effector prediction, homology-based inference implies finding a sequence-similar experimentally annotated type III effector (‘Methods’ section), *e.g.* via PSI-BLAST.

The accuracy of homology-based inference by PSI-BLAST was comparable to that of our *de novo* prediction method on the cross-validation *Development set* (Table 1: 91% vs. 92%). However, at this level of accuracy, its coverage was significantly higher (Table 1: 84% vs. 60%). This result encouraged combining these two approaches as introduced in our recent work, LocTree3^23^: use PSI-BLAST when sequence similarity suffices (*e*-value≤10^−3^; Table 1: F_1_ = 0.87 complete set) and the SVM otherwise (Table 1: F_1_ = 0.67 on subset of proteins without PSI-BLAST hit). The combined method, pEffect, outperformed both its components, reaching an F_1_ measure of 0.91 (Table 1).

**Table 1.**
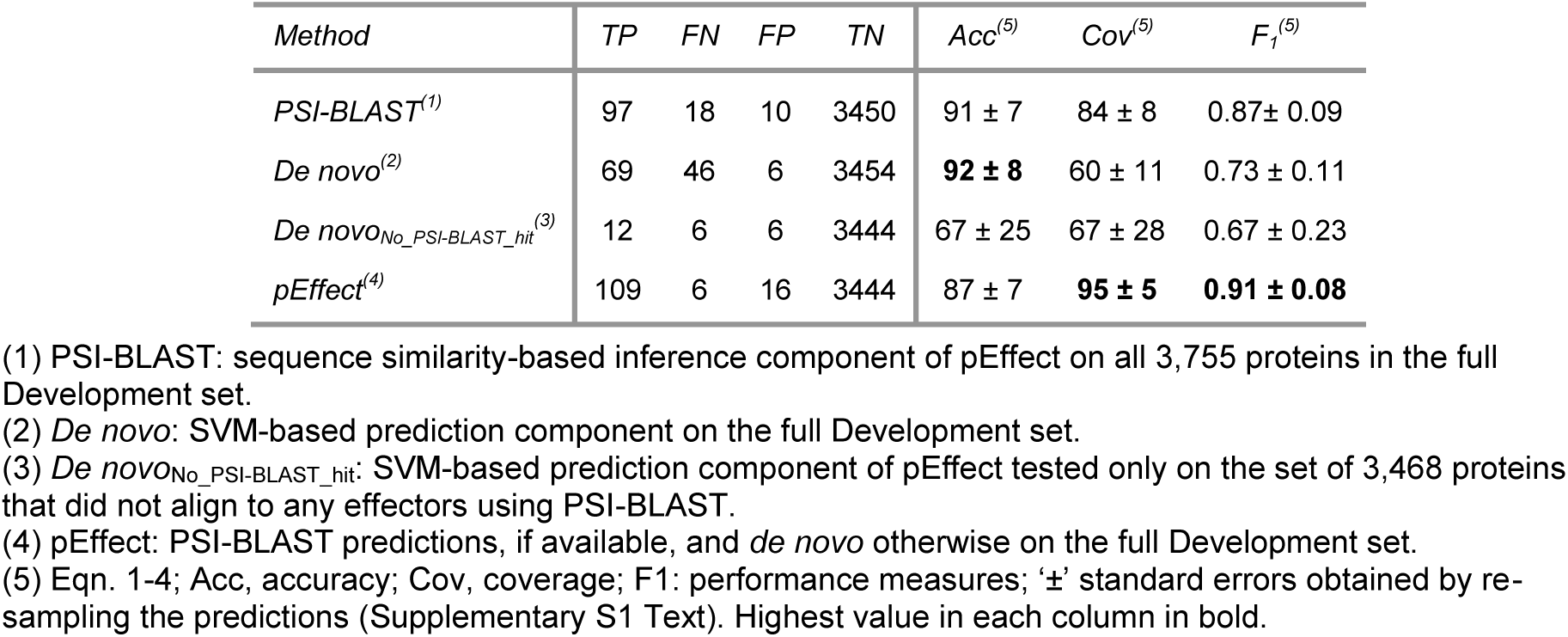
Performance of pEffect and its components on the Development set.

| Method | TP | FN | FP | TN | Acc<sup>(5)</sup> | Cov<sup>(5)</sup> | F<sub>1</sub><sup>(5)</sup> |
| --- | --- | --- | --- | --- | --- | --- | --- |
| PSI-BLAST<sup>(1)</sup> | 97 | 18 | 10 | 3450 | 91 ± 7 | 84 ± 8 | 0.87 ± 0.09 |
| De novo<sup>(2)</sup> | 69 | 46 | 6 | 3454 | **92 ± 8** | 60 ± 11 | 0.73 ± 0.11 |
| De novo<sub>No_PSI-BLAST_hit</sub><sup>(3)</sup> | 12 | 6 | 6 | 3444 | 67 ± 25 | 67 ± 28 | 0.67 ± 0.23 |
| pEffect<sup>(4)</sup> | 109 | 6 | 16 | 3444 | 87 ± 7 | **95 ± 5** | **0.91 ± 0.08** |

(1) PSI-BLAST: sequence similarity-based inference component of pEffect on all 3,755 proteins in the full Development set.
(2) De novo: SVM-based prediction component on the full Development set.
(3) De novo<sub>No\_PSI-BLAST\_hit</sub>: SVM-based prediction component of pEffect tested only on the set of 3,468 proteins that did not align to any effectors using PSI-BLAST.
(4) pEffect: PSI-BLAST predictions, if available, and de novo otherwise on the full Development set.
(5) Eqn. 1-4; Acc, accuracy; Cov, coverage; F<sub>1</sub>: performance measures; '±' standard errors obtained by re-sampling the predictions (Supplementary S1 Text). Highest value in each column in bold.

### pEffect outperformed other methods

We compared pEffect to publicly available methods: BPBAac^13^, EffectiveT3^12^, T3_MM^24^, Modlab^15^ and BEAN 2.0^14^. BPBAac, EffectiveT3, T3_MM and Modlab focus exclusively on N-terminal features, while BEAN 2.0 and pEffect are not confined to those regions only (Methods, Supplementary S2 Text). BPBAac, T3_MM and Modlab rely solely on amino acid composition; EffectiveT3 combines amino acid composition and secondary structure information; BEAN 2.0 uses BLAST^18^ and PFAM^25^ domain searches, evolutionary information encoded in N-and C-termini, as well as information from an intermediate sequence region. We compared performance for UniProt^26^ proteins that had NOT been used to develop any method, and for T3DB^11^ proteins, some of which all methods (incl. pEffect) had used for development. In our hands, pEffect significantly outperformed its competitors on UniProt sets containing eukaryotic proteins (Fig. 1, Supplementary Table S1). The F_1_ performance of pEffect exceeded the other methods by more than 0.35 (∆F_1_ = (pEffect, BEAN 2.0) = 0.35 for both UniProt sets, Supplementary Table S1). On the bacterial T3DB data sets, pEffect performed within one standard error of the prediction performance achieved by its best performing competitor BEAN 2.0. Thus, pEffect performed as well or better when benchmarked with existing tools in distinguishing type III effectors from bacteria (F_1_>0.64*) and from eukaryotes* (F_1_>0.88). This improvement is particularly important to, *e.g.* annotate results from contaminated metagenomic studies^27^.

**Figure 1:**
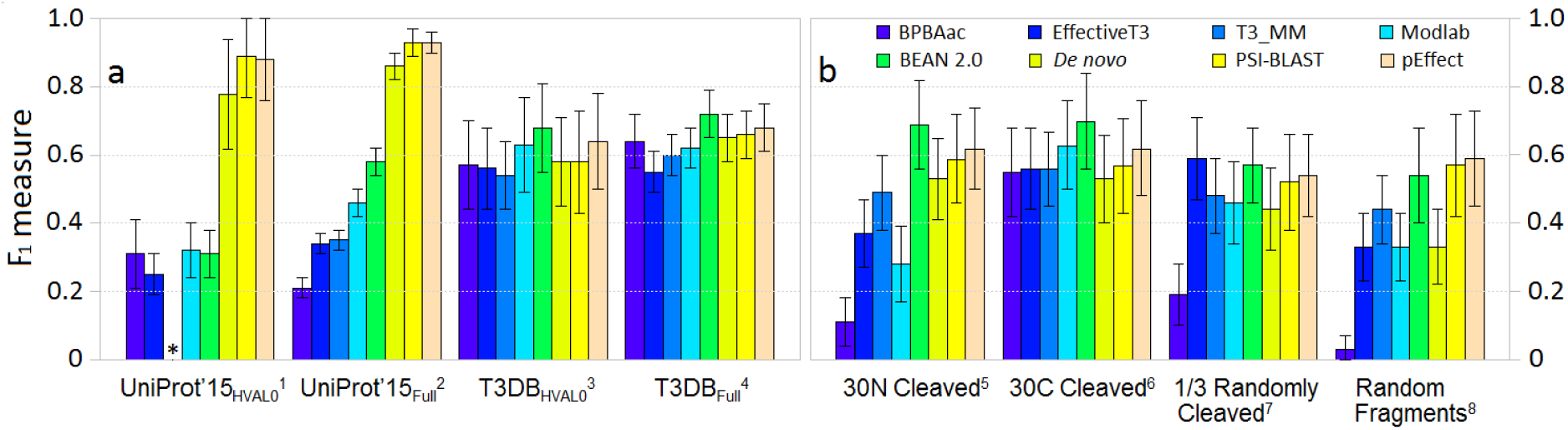
Method performance comparison on independent test sets and protein fragments. Performance (Supplementary S1 Text: F_1_ measure, Eqn. 3; ‘±’ standard error, Eqn. 4) was measured for BPBAac, EffectiveT3, T3_MM, Modlab and BEAN 2.0 methods (Supplementary S2 Text). We also computed F_1_ for *de novo* (SVM-based) predictions alone, PSI-BLAST homology-based look up alone, and pEffect: a combination of PSI-BLAST (if a hit is available) and *de novo* (otherwise). **Panel (a)** shows performance on evaluation data sets (Methods) including ^**(1)**^UniProt’15_HVAL0_ (51 effectors and 691 non-effector bacterial and eukaryotic proteins, added to UniProt after 2014_02 release, sequence homology reduced at HVAL<0), ^**(2)**^UniProt’15_Full_ (498 effectors and 1,509 non-effector bacterial and eukaryotic proteins added to UniProt after 2014_08 release, NOT homology reduced), ^**(3)**^T3DB_HVAL0_ (66 effectors and 128 non-effector bacterial proteins from T3DB database, sequence homology reduced at HVAL<0), and ^**(4)**^T3DB_Full_ (218 effectors and 831 non-effector bacterial proteins from T3DB database, NOT homology reduced). Note: T3_MM was not able to produce results for the UniProt’15_HVAL0_ set during manuscript preparation. **Panel (b)** shows performance on *fragments* produced from ^**(3)**^T3DB_HVAL0_ (Methods) including ^**(5)**^approach i: 30 N-terminal amino acids cleaved off, ^**(6)**^ii: 30 C-terminal amino acids cleaved off, ^**(7)**^iii: Randomly selected two thirds of the protein sequence, and ^**(8)**^iv: Randomly selected sequence fragments of typical translated read length (average 110 amino acids, Supplementary Fig. S1).

### pEffect excelled even for protein fragments

To evaluate pEffect’s ability to annotate effectors from incomplete genomic assemblies and mistakes, we fragmented the proteins from the homology reduced T3DB_HVAL0_ set containing bacterial proteins only. We started with protein rather than gene sequence fragments, because we did not expect incorrect gene translations of DNA reads, even if sufficiently long, to trigger incorrect effector predictions from any method. Four different approaches were used to generate protein fragments: (i) remove the first 30 residues (N-terminus) from the full protein sequence, (ii) remove the last 30 residues (C-terminus), (iii) randomly remove one third of residues, and (iv) randomly choose from each protein a single fragment of a typical translated read length (Supplementary Fig. S1).

pEffect compared favourably to all other methods for all fragment sets (i-iv). Most methods performed best on fragments with C-terminal cleavage (set ii, Fig. 1, Supplementary Table S2). Performance was lowest for random fragments of typical read lengths (set iv). For pEffect it dipped least (F_1_ = 0.59±0.14 on set iv *vs*. F_1_ = 0.64±0.14 on full length, Supplementary Table S1). For all fragment sets, performances of homology-based approaches, *i.e*. of PSI-BLAST, pEffect and BEAN 2.0 were within the standard error of the performance obtained when using full-length sequences (T3DB_HVAL0_ set; Fig. 1, Supplementary Table S1). These results suggest that the features distinguishing type III effectors are spread over the entire protein sequence and can be picked up by local alignment.

### Reliability index identified confident predictions

pEffect provides a reliability index (RI) to measure the confidence of a prediction; the value of RI ranges from 0 (uncertain) to 100 (most reliable). For PSI-BLAST searches, RIs are normalized values of percentage pairwise sequence identities read of the alignments. For *de novo* predictions, RIs are values corresponding to SVM scores (Methods). Including predictions with low RIs gives many trusted results at reduced accuracy. Higher accuracy predictions are obtained by sampling at higher RIs, thus reducing the total number of trusted samples. For example, at the threshold of RI≥50, over 87% of all predictions of type III effectors are correct and 95% of all effectors in our set are identified (Supplementary Fig. S2: black arrow). On the other hand, at RI>80 effector predictions are correct 96% of the time, but only 78% of all effectors in the set are identified (Supplementary Fig. S2: gray arrow). Thus, users can choose the most appropriate threshold for a given study. Users can also focus on previously unidentified effectors (*de novo* predictions) or, *vice versa*, on validated homologs of known effectors (PSI-BLAST matches; Supplementary Fig. S3).

### Application of pEffect: scanning proteomes for type III effectors

We used pEffect to annotate type III effectors in 862 bacterial (274 Gram-positive, 588 Gram-negative bacteria) and 90 archaeal proteomes from the European Bioinformatics Institute (EBI: http://www.ebi.ac.uk/genomes/; predictions available on the pEffect website). Each bacterium was predicted to have some type III effectors, with a minimum of 0.8% of the proteome −2 out of all 240 proteins – identified as effectors (Fig. 2, Supplementary Table S3). For some Gram-negative bacteria, over 750 type III effectors were predicted (Supplementary Table S3), *e.g*. 870 effectors in *S. aurantiaca* DW4/3-1, which is indeed known to have a T3SS and effectors^28^.

**Figure 2:**
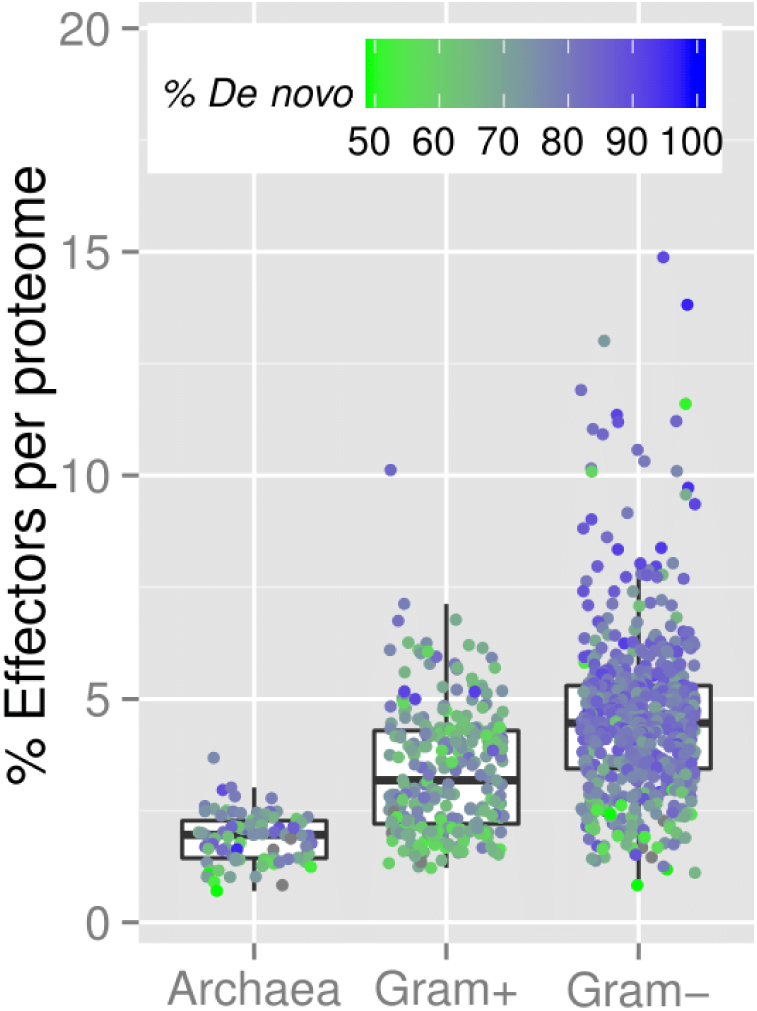
Percentage of predicted effectors in full proteomes. The figure shows the box-plot-and-instance representation of percentages of pEffect-predicted type III effectors (Y-axis) in 90 archaeal, 274 Gram-positive and 588 Gram-negative bacterial organisms (X-axis), which are shown as dots. At least 50% of effector predictions in all, except 11 organisms in our set were predicted *de novo*. In the figure, the colour represents the percentage of *de novo* predictions for each organism: from green (50% *de novo,* 50% PSI-BLAST) to blue (100% *de novo*, 0% PSI-BLAST). While effectors predicted in archaea and Gram-positive bacteria are often picked up by PSI-BLAST, effectors in Gram-negative bacteria are mostly *de novo* predictions (mostly blue dots).

Overall, the number of predicted type III effectors was 1-10% of the whole proteome in Gram-positive bacteria and 1-15% in Gram-negative bacteria (Fig. 2, Supplementary Table S3). To further understand our predictions, we retrieved UniProt keywords of predicted effectors. Their annotations varied widely, with the most common for both types of bacteria being *transferase*, depicting a large class of enzymes that are responsible for the transfer of specific functional groups from one molecule to another, *nucleotide-binding* – a common functionality of effector proteins, *ATP-binding* – an essential component of T3SS, and *kinase* – necessary for the expression of T3SS genes. About one fourth (26-29% per proteomes) of predicted type III effectors are functionally ‘unknown’ (Supplementary Table S4).

We also predicted type III effectors in all archaeal proteomes, with over 100 effectors identified in the proteomes of *H. turkmenica* DSM 5511 and *M. acetivorans* C2A (126 and 105 effectors, respectively; Supplementary Table S3). On average, there were fewer effectors predicted in archaea than in bacteria: 1.9% is the overall per-organism number for archaea *vs*. 3.4% for Gram-positive and 4.6% Gram-negative bacteria (Fig. 2). The most frequent annotations of predicted archaeal effectors were similar to those for predicted bacterial effectors, namely ‘unknown’, nucleotide-binding, ATP-binding and transferase (Supplementary Table S4).

### Evaluation of predictions for proteomes

We BLASTed proteins representative of five T3DB Ortholog clusters (*e*-value≤10^−3^; Supplementary Table S5) against the full proteomes of our 862 bacteria and 90 archaea set. We thus aimed to identify those proteomes likely equipped with the T3SS machinery (Fig. 3).

**Figure 3:**
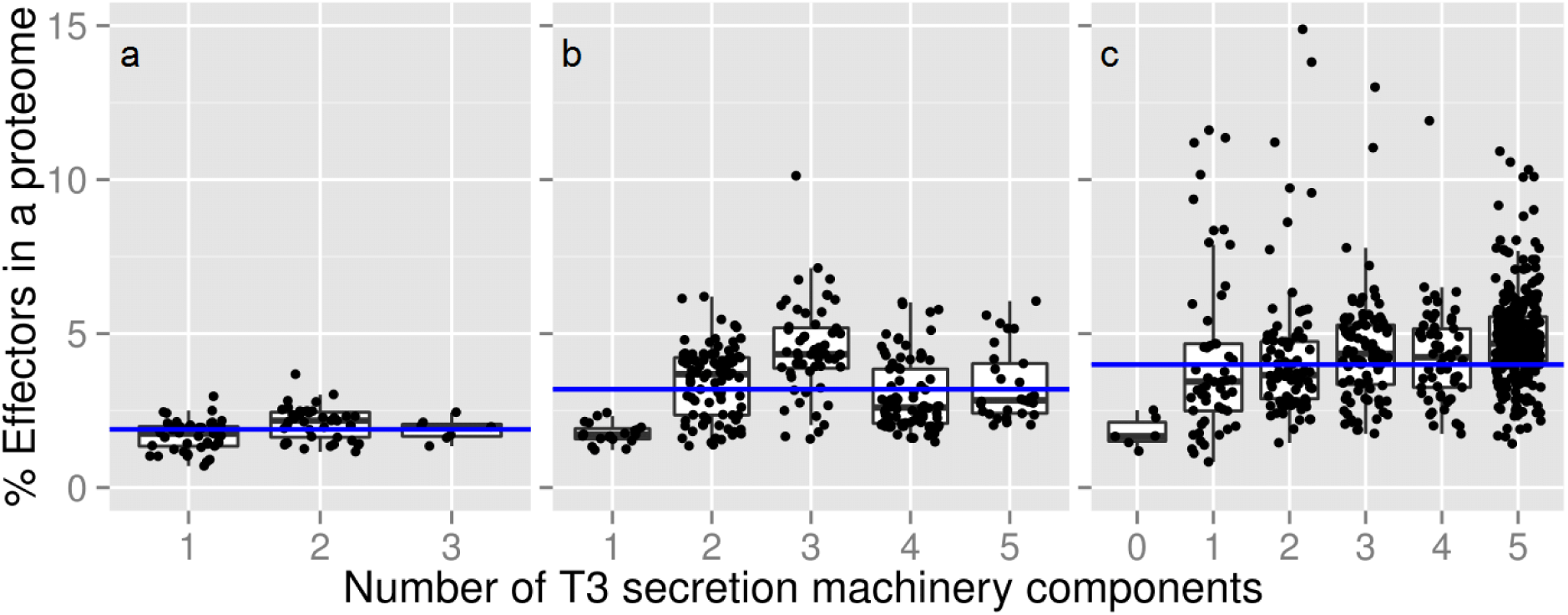
Proteomes encoding some of the five components of T3SS machinery. **(a)** 90 archaea proteomes, **(b)** 274 Gram-positive bacteria and **(c)** 588 Gram-negative bacteria were scanned for the presence of T3SS and are shown as dots in the figure. The percentage of type III effectors predicted by pEffect (Y-axis) is compared to the number of type III secretion machinery components (max. five T3 Ortholog clusters; Methods) identified in these proteomes (X-axis). Note that effector predictions are computationally completely independent of machinery component identifications. While type III effectors compose up to 3.7% of an archaeal proteome (mean 1.9%, blue horizontal line), this number is much larger for bacteria, reaching up to 10.1% of an entire proteome for Gram-positive bacteria (mean 3.4%), and 14.9% for Gram-negative bacteria (mean 4.6%; for those with five T3SS components, mean 4.8%). Note that six Gram-negative bacterial species did not contain detectable homologs of any of the required machinery components (not even ATPases), indicating that their genomes are further diverged than those of other species.

We found that, as expected, archaea never contain a full T3SS (maximum three out of five components). In Gram-negative bacteria, the number of predicted effectors correlated much better with the number of type III machinery components (Pearson correlation *r* = 0.37) than in Gram-positive bacteria (*r* = 0.13). The combination of a high percentage of predicted type III effectors and a high number of conserved type III machinery components provides strong evidence for the presence of the type III secretion abilities (Fig. 3). As a rule of thumb, based on our observations in archaea and Gram-positive bacteria, we suggest that these abilities can be reliably identified by the presence of the complete T3SS and ≥5% of the genome dedicated to effectors. With these cutoffs, 20% (115 species) of the Gram-negative bacteria in our set are identified as type III secreting. We randomly picked ten species from these 20% and found evidence in the literature for T3SS presence for seven of them (Supplementary Table S6). No archaeal species and only five Gram-positive bacteria fit these cutoffs. Note that our rule does not imply that organisms with full T3SS and over 5% predicted effectors necessarily have the complete ability to use the system. Instead, we suggest that organisms without the necessary components cannot use the system. Overall, our results indicate that the experimental annotation of the type III secretion in isolated and cultured organisms is incomplete, leaving significant room for improvement, possibly with assistance from pEffect.

Finally, we extracted from the HAMAP database^29^ available annotations of pathogenicity and symbiotic relationships for 115 Gram-negative bacteria in our set with a complete T3SS and ≥5% of the genome dedicated to effectors. We compared the number of predicted effectors in organisms that infect eukaryotic cells in general and mammalian cells in particular with those that are currently not known to be symbiotic or pathogenic. Note that further manual curation of currently not annotated bacteria still highlights possibility of type III secretion for a large fraction of them^30–32^. Our analysis showed that while the distributions of numbers of effectors across the different types of bacteria were not significantly different, mammalian pathogens carried, on average, more effectors than pathogens of other taxa. Those, in turn, carried more effectors than bacteria not currently annotated as pathogenic or symbiotic (Supplementary Fig. S4). Thus, we believe that pEffect can be used to pinpoint for future exploration of the type III secretion-mediated pathogenicity of newly sequenced organisms.

## Discussion

pEffect successfully combines complementary approaches for the prediction of type III effector proteins: homology-based and *de novo*. Specifically, it uses PSI-BLAST for a high accuracy (precision) mode of prediction and SVM for improved coverage (recall). The resulting single method pEffect outperforms both of its individual components (Table 1) and other methods (Fig. 1, Supplementary Tables S1-S2). When tested on samples contaminated with eukaryotic proteins, pEffect predicts effectors with a performance level that is significantly higher than that of any other currently available method (Fig. 1, Supplementary Tables S1 and S7). Similar to the results of Arnold *et al.^12^*, we find that there is no significant difference in performance across different species of bacteria (pEffect: F1= 0.54±0.31on a data set with no proteins of the same species shared between training and test sets vs. F1= 0.91±0.08 on the Development set). pEffect was trained on a sequence homology reduced data set at HVAL=0 (*i.e*. there is no pair of sequences in our data set with over 20% sequence identity that have over 250 amino acid residues aligned) that to our knowledge presents the largest and most complete set of effector proteins currently available. The data set can be downloaded from pEffect’s website.

pEffect uses information stored in the entire protein sequence and performs on sequence fragments just as well as on full-length protein sequences (Fig. 1, Supplementary Table S2). This result made us conclude that signals discriminating effector proteins are distributed across the entire protein sequences and are not confined to the N-terminus, as it is currently anticipated. This finding was surprising and extremely relevant for the analysis of metagenomic read data. Deep Sequencing (or NGS) produces immense amounts of DNA reads, which need to be assembled and annotated to be useful. Erroneous (chimeric) gene assemblies or wrong gene predictions are common in sequencing projects^33^. To bypass the assembly errors when identifying type III secretion activity in a particular metagenomic sample it would help to annotate effectors from peptides translated directly from the DNA reads. pEffect facilitates this type of direct analysis of metagenomic sequence data, establishing the level of type III secretion activity and, by proxy, the endosymbiotic interactions and the potential presence or absence of pathogenic organisms in a particular environment.

We applied pEffect to over 900 prokaryotic proteomes with the aim of annotating those organisms that are likely to utilize a T3SS. We validated our results using three different metrics: (i) percentage of predicted effector proteins per proteome, (ii) evolutionary age of an organism and (iii) the number of conserved T3SS elements. As expected, pEffect predicted a higher percentage of effector proteins per proteome in Gram-negative bacteria with full T3SS (five conserved T3SS elements) than in Gram-positive bacteria and archaea that are not known to utilize the system (Fig. 2–3). This indicates a possible acquisition of a larger effector repertoire in Gram-negative bacteria, which was unnecessary for other organisms. Incorporating the independently established evolutionary age estimate, effector proteins of T3SS-using Gram-negative bacteria appear to further diversify with the increasing evolutionary distance from the last common ancestor (Fig. 4a). This correlation could not be expected at random, as the age of bacteria and their effector quantities are independently established and are not correlating for other organisms.

Interestingly, homology searches have identified roughly equal numbers of effectors (on average, 1% of each respective proteome; Supplementary Table S3) across both types of bacteria. As their percentage per proteome remains stable over time (Supplementary Table S3) and as they are found in almost all organisms with PSI-BLAST, we suggest these effectors to be the older ones that had the time to spread throughout different species. On the other hand, the increasing number of new effectors, recognized by the SVM, in relationship to organism age (as long as organism is using T3SS, Fig. 4b), indicates likely new “inventions” that accumulate over time of T3SS use. These results are in line with potential ancestral presence of the early complete secretory system^10,34^, including the machinery and the secreted proteins, and further diversification of effectors exclusively in T3SS-utilizing Gram-negative bacteria.

**Figure 4:**
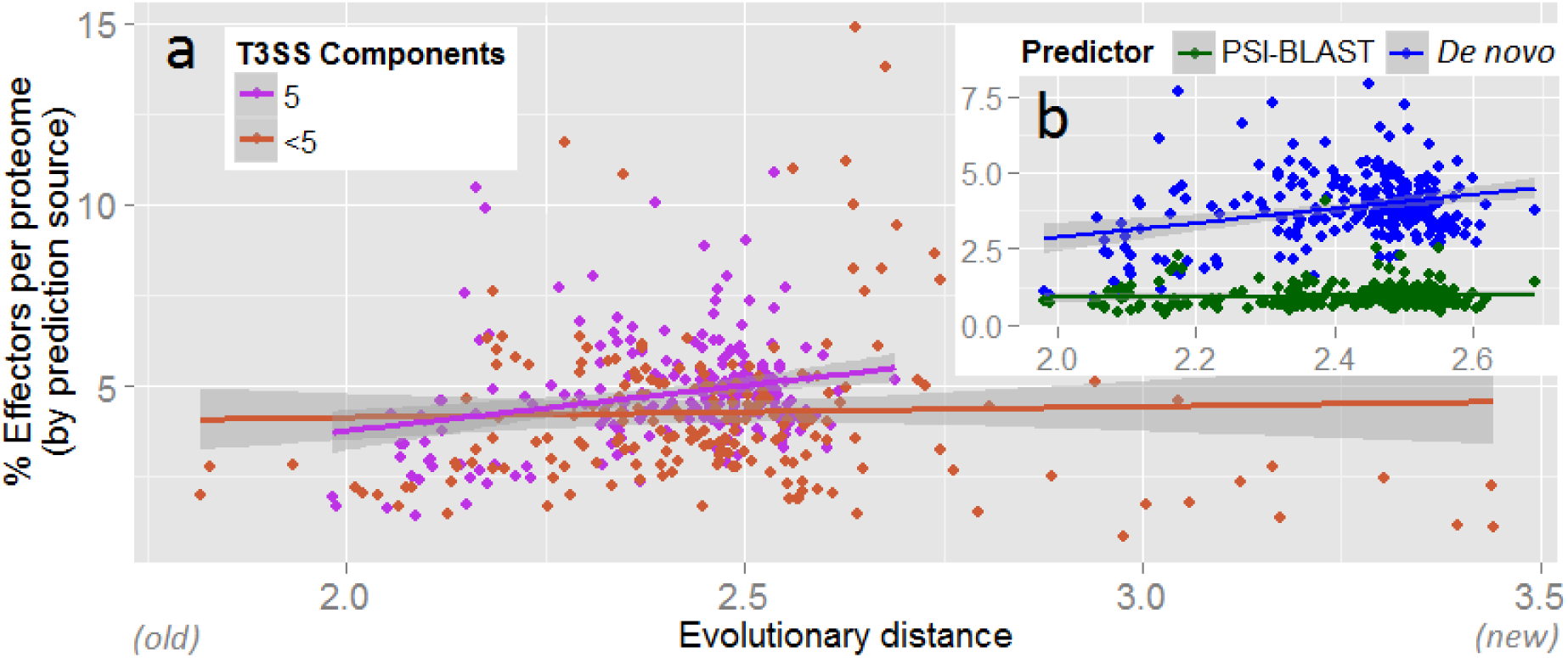
pEffect’s whole proteome predictions in Gram-negative bacteria. **(a)** pEffect predicted type III effector proteins in the proteomes of 294 Gram-negative bacteria. The proteomes are shown as red and purple dots. Purple dots indicate proteomes with five type III machinery components (full T3SS) and red dots are proteomes with fewer components. For each proteome, the evolutionary distance from the last common ancestor (X-axis), extracted from Lang *et al*.^35^, is plotted against the percentage of proteins predicted as effectors (Y-axis). While there is a correlation between the age and the quantities of effectors in proteomes of organisms with full T3SS (purple trend-line), the same appears not to be the case for organisms with less than five components. **(b)** Proteomes with full T3SS identified by source. Green dots are the percentage of proteins predicted as effectors by homology searches (PSI-BLAST) and blue dots are *de novo* predictions. While PSI-BLAST appears to consistently pick up ~1% of each proteome of all organisms (green horizontal trend-line), the effectors in Gram-negative bacteria diversify further over evolutionary distance, as indicated by the increase in the number of *de novo* predictions.

The set of *de novo-*identified effectors found across bacteria is a good starting point for further investigation into effector origins. Due to T3SS significance in pathogenicity of Gram-negative bacteria, the *de novo* identified effectors are also potentially interesting as drug targets.

pEffect’s high prediction accuracy raises an interesting question about its false positive predictions of effectors in Gram-positive bacteria, which is not known to utilize T3SS. Roughly one fourth of these predicted effectors are of yet-unknown function. Those that are annotated include enzymes necessary for flagellar motility (Supplementary Table S6). This finding is in line with evidence of shared ancestry between bacterial flagellar and type III secretion systems^9^. Gene genealogies^20^ and protein network analysis approaches^21^ suggest independent evolution of both systems from a common ancestor, comprising a set of proteins forming a membrane-bound complex. The fact that the flagellar system can also secrete proteins^36^ suggests that this ancestor may have played a secretory role^9^. The pervasiveness of the flagellar apparatus across the bacterial space also suggests that the ancestral complex existed prior to the split of the cell-walled and double-membrane organisms, indicated by the differences in gram staining. Thus, it is not surprising that we find T3SS component homology in Gram-positive bacteria even in the absence of type III secretion functionality. Curiously, our results show that the loss of the type III secretion functionality, indicated by the loss of the complete T3SS, has proceeded at a roughly similar rate in Gram-positive and Gram-negative bacteria (Fig. 5a); *i.e*. once the T3SS becomes incomplete (4 components) and, arguably, non-functional, further loss of components consistently follows. Notably, a complete T3SS is only visible in early Gram-positive bacteria, but preserved across time in Gram-negative bacteria (Fig. 5b), further confirming the likely presence of the ancestral secretory complex in the last common bacterial ancestor.

**Figure 5:**
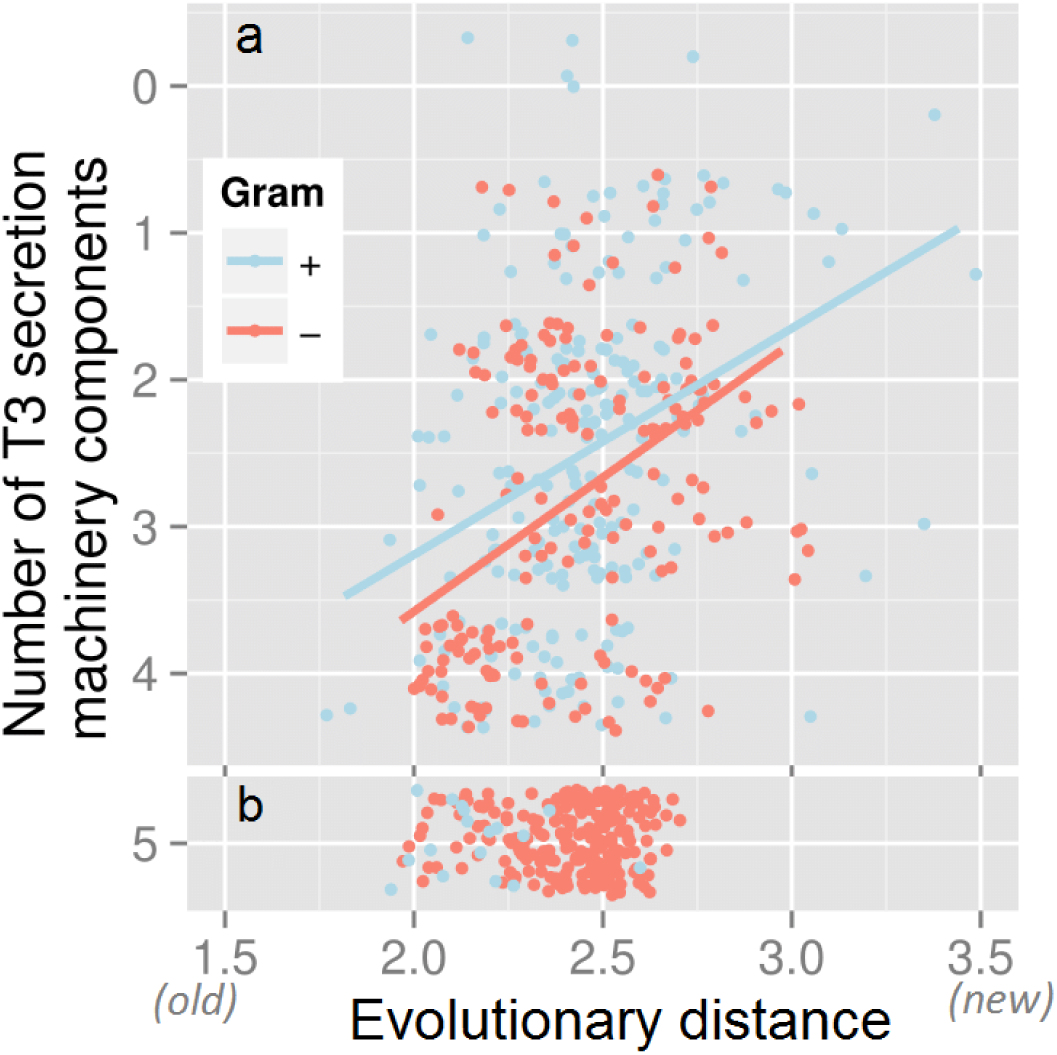
Loss of T3SS functionality differentiates Gram-positive and -negative bacteria. 274 Gram-positive bacteria (blue dots) and 588 Gram-negative bacteria (red dots) are screened for the number of conserved components of T3SS (max. 5 T3DB Ortholog clusters; Material and Methods) in their genomes (Y-axis) and plotted against the evolutionary distance from the most recent common ancestor (X-axis). Once the T3SS is lost **(a)**, *i.e*. less than 5 components are present, further rate of loss of components is the same for all bacteria. The number of Gram-negative bacteria with the complete system **(b)**, *i.e*. all 5 components present, however, remains constant across evolutionary time, while the number of Gram-positive bacteria declines.

pEffect also predicts a significant number of false positive effectors in archaea, inspiring the question: did T3SS exist before the archaea/bacteria split? Unfortunately, the presence of the beginnings of T3SS in the common ancestor of bacteria and archaea is neither directly supported nor negated by our results. Archaeal flagella have little or no structural similarities to bacterial flagella and none of the archaea that we tested had the complete T3SS (Fig. 2). If the common ancestor of archaea and bacteria did encode the core ancestral complex, the latter observation would indicate a loss of functionality in archaea. Another possibility is that the T3SS in bacteria may have been built over time from duplicated and diversified paralogous genes of the core complex after the archaea/bacteria split. In both of these scenarios, the prediction of type III effectors in archaea would indicate re-purposing of the proteins secreted by the ancestral complex. In fact, 0.5% of an average archaeal genome is identified by homology to known effectors and another 0.9% *de novo-*identified proteins are homologous (PSI-BLAST *e*-value≤10^−3^) to *de novo*-identified effectors of Gram-negative bacteria. These proteins must have been re-purposed in modern archaea; in fact, they are annotated with a range of molecular functionalities (Supplementary Table S6). The use of an additional 0.5% of the archaeal proteome that is picked up by pEffect *de novo* and has no homologs in bacteria remains an enigma. While similarity between archaeal proteins and bacterial type III effectors and machinery is insufficient to draw definitive conclusions regarding common ancestry, it is significant for further exploration; *i.e*. if roughly one tenth of the identified effectors of Gram-negative bacteria and half of the machinery have homologs in archaea, could there have been a common ancestral secretion complex that has developed early on in evolutionary time and has given root to many systems observed today?

pEffect immediately and importantly contributes to the study of type III secretion mechanisms. It allows for rapid identification of type III secretion abilities within unassembled genomic and metagenomic read data. Moreover, the quantity of identified effectors seems to correspond with bacterial pathogenicity, potentially contributing to the tracking of infectious strains. We believe that pEffect will facilitate future experimental insights in microbiological research and will significantly contribute to our understanding and management of infectious disease.

## Methods

### Development data sets

Our positive data set of known type III effector proteins was extracted from scientific publications^12,37−44^ and the Pseudomonas-Plant Interaction web site (http://www.pseudomonas-syringae.org/). We additionally queried Swiss-Prot with keywords ‘type III effector’, ‘type three effector’ and ‘T3SS effector’ and manually curated the results for experimentally identified effectors (removing entries with “potential”, “probable” and “by similarity” annotations). All corresponding amino acid sequences were taken from UniProt^26^, 2012_01 release. In total, our positive (effector) data set contained 1,388 proteins.

To compile our negative data set of non-type III effectors we used the experimentally annotated Swiss-Prot proteins from the 2012_01 UniProt release. We extracted all bacterial proteins that were NOT annotated as type III effectors and had no significant sequence similarity (BLAST *e*-value>10) to any type III effector in our positive set. We also added all eukaryotic proteins applying no sequence similarity filters. Our negative set thus contained roughly 470,000 proteins.

We removed from our sets all proteins that were annotated as ‘uncharacterized’, ‘putative’, or ‘fragment’. We reduced sequence redundancy independently in each set using UniqueProt^45^, ascertaining that no pair of proteins in one set had alignment length of less than 35 residues or a positive HSSP-value^46,47^ (HVAL≥0). After redundancy reduction our sequence-unique sets contained 115 type III effector proteins from 43 different bacterial species and 3,460 non-effector proteins (of which 37% were bacterial). Note that proteins from positive and negative sets were sometimes similar as homology reduction was only applied *within* sets and not *across* sets. Here, this set of sequences (positive and negative sets together) is termed the *Development set*. All pEffect performance results were compiled on stratified cross-validation of this Development set (five-fold cross-validation, *i.e*. we split the entire set into five similarly-sized subsets and trained five models, each on a different combination of four of these subsets, testing each model on every subset exactly once).

### Additional data sets

Comparing pEffect performance to that of other methods using our cross-validation approach has only limited value due to the possible overlap between our testing and other methods’ training sets, and can lead to an overestimate of other methods’ performance. A more meaningful way is to use non-redundant sets of effector and non-effector proteins that have never been used for the development of any method. Toward this end, we extracted the following data sets:

(1) We collected all type III effectors added to UniProt after the 2014_02 release and non-type III bacterial and eukaryotic proteins added to Swiss-Prot after the same release. These were redundancy reduced at HVAL<0 to produce the UniProt’15_HVAL0_ set (51 effectors and 691 non-effectors, of which 53% were of bacterial origin). Note that additionally reducing this set to be sequence dissimilar to the *Development set* would retain only 10 type III effectors, too few for reliable performance estimates. However, even for this smaller and completely independent set, the performance of pEffect was higher than of other tools, making pEffect a uniquely reliable method for determining new effectors (Supplementary Table S7).

(2) To answer the question “how well will pEffect perform on protein sequences added to databases within the next six months?” we collected the proteins added to UniProt (type III effectors) and Swiss-Prot (non-effector bacterial and eukaryotic sequences) after the 2014_08 release, producing the set *UniProt’15*_*Full*_ (498 effectors and 1,509 non-effectors, of which).

(3) We also extracted all bacterial type III effectors from the T3DB database^11^ – *T3DB*_*Full*_ set (218 effectors and 831 non-effectors). We deliberately kept the redundancy in this set (up to HVAL = 66, *i.e*. over 85% pairwise sequence identity over 450 residues aligned). Note that some proteins from this set are contained in the training sets of all compared methods, including pEffect.

(4) Finally, we redundancy reduced T3DB set at HVAL<0. This gave the *T3DB*_*HVAL0*_ set (66 effectors and 128 non-effectors).

### T3DB Ortholog clusters of the type III secretion system (T3SS) machinery

T3DB is a database of experimentally annotated T3SS-related proteins in 36 bacterial taxa. Proteins of the same function and the same evolutionary origin are clustered in T3DB into *T3 Ortholog* clusters (http://biocomputer.bio.cuhk.edu.hk/T3DB/T3-ortholog-clusters.php). The proteins of these clusters form ten components of the T3SS. Proteins of five of these components (export apparatus, inner membrane ring, outer membrane ring, cytoplasmic ring, and ATPase) are present in all 36 taxa in T3DB (Supplementary Table S2). We thus defined the minimum number of five components necessary for the formation of the T3SS machinery. With the exception of the outer membrane ring, these components have also been defined as the core before^9^.

### Prediction methods

We tested several ideas for prediction, including the following:

#### Homology-based inference

We transferred type III effector annotations by homology using PSI-BLAST^18^ alignments. For every query sequence we generated a PSI-BLAST profile (two iterations, inclusion threshold *e*-value≤10^−3^) using an 80% non-redundant database combining UniProt^48^ and PDB^49^. We then aligned this profile (inclusion *e*-value≤10^−3^) against all type III effectors extracted from the literature and the UniProt 2012_01 release. For known effectors, we excluded the PSI-BLAST self-hits. We transferred annotation to the query protein from the hit with highest pairwise sequence identity of all retrieved alignments.

#### De novo prediction

We used the WEKA^50^ Support Vector Machine (SVM)^19^ implementation to discriminate between type III effector and non-effector proteins. For each protein sequence, we created a PSI-BLAST profile (as described above) and applied the Profile Kernel function^51,52^ to map the profile to a vector indexed by all possible subsequences of length *k* from the alphabet of amino acids; we found that *k* = 4 amino acids provides best results. Each element in the vector represents one particular *k*-mer and its score gives the number of occurrences of this *k*-mer that is below a certain user-defined threshold σ; we found that σ = 7 provides best results. This score is calculated as the ungapped cumulative substitution score in the corresponding sequence profile. Thus, the dot product between two *k*-mer vectors reflects the similarity of two protein sequence profiles. Essentially, the method identifies those stretches of *k* adjacent residues in profiles of type III effectors that are most informative for prediction and matches these to the profile of a query protein. The parameters for the SVM and the kernel function were determined separately for each fold in our 5-fold cross-validation and, thus, were never optimized for the test sets.

#### pEffect

Our final method, pEffect, combined sequence similarity-based and *de novo* predictions. Toward this end, over-fitting was avoided through the simplest possible combination: if any known type III effector is sequence similar to the query use this (similarity-based prediction), otherwise use the *de novo* prediction.

#### Reliability index

The strength of a pEffect prediction is represented by a reliability index (RI) ranging from 0 (weak prediction) to 100 (strong prediction). For *de novo* predictions, we computed RI by multiplying the SVM output by 100 for positive (type III effector) predictions and subtracted this score from 100 for negative predictions. For sequence similarity-based inferences, the RI is the percentage of pairwise sequence identity normalized to the interval [50, 100], to agree with the SVM prediction range.

### Evolutionary distances

For the discovery of novel type III effectors in entirely sequence organisms, we extracted evolutionary distances from the phylogenetic tree of 2,966 bacterial and archaeal taxa, inferred from 38 concatenated genes and available in the Newick format^35^.

## Acknowledgements

Thanks to Tim Karl, Guy Yachdav, Laszlo Kajan (all TUM) and Yannick Mahlich (Rutgers) for invaluable help with hardware and software; to Chengsheng Zhu (Rutgers) for helpful discussions; to Jessie Maguire (Rutgers), Marlena Drabik, Inga Weise and Lothar Richter (all TUM) for administrative support. Last, not least, thanks to all those who deposit their experimental data in public databases, and to those who maintain these databases.

## Author Contributions Statement

T.G. and Y.B. designed the project. T.G. developed the prediction method, its web server, and performed whole proteome predictions. T.G., Y.B and B.R. analysed the results and wrote the manuscript.

## Additional Information

Competing financial interests: The authors declare no conflicts of interest.

## Supplementary Information

Supplementary information is available at the Nature Scientific Reports website.

## References

1 Holland, I. B., Schmitt, L. & Young, J. Type 1 protein secretion in bacteria, the ABC-transporter dependent pathway (review). Molecular membrane biology 22, 29–39 (2005).

2 Nivaskumar, M. & Francetic, O. Type II secretion system: a magic beanstalk or a protein escalator. Biochimica et biophysica acta 1843, 1568–1577, doi:10.1016/j.bbamcr.2013.12.020 (2014).

3 Cornelis, G. R. The type III secretion injectisome. Nature reviews. Microbiology 4, 811–825, doi:10.1038/nrmicro1526 (2006).

4 Low, H. H. et al. Structure of a type IV secretion system. Nature 508, 550–553, doi:10.1038/nature13081 (2014).

5 Leo, J. C., Grin, I. & Linke, D. Type V secretion: mechanism(s) of autotransport through the bacterial outer membrane. Philosophical transactions of the Royal Society of London. Series B, Biological sciences 367, 1088–1101, doi:10.1098/rstb.2011.0208 (2012).

6 Russell, A. B., Peterson, S. B. & Mougous, J. D. Type VI secretion system effectors: poisons with a purpose. Nature reviews. Microbiology 12, 137–148, doi:10.1038/nrmicro3185 (2014).

7 Hueck, C. J. Type III protein secretion systems in bacterial pathogens of animals and plants. Microbiology and molecular biology reviews : MMBR 62, 379–433 (1998).

8 Troisfontaines, P. & Cornelis, G. R. Type III secretion: more systems than you think. Physiology 20, 326–339, doi:10.1152/physiol.00011.2005 (2005).

9 McCann, H. C. & Guttman, D. S. Evolution of the type III secretion system and its effectors in plant-microbe interactions. The New phytologist 177, 33–47, doi:10.1111/j.1469-8137.2007.02293.x (2008).

10 Figueira, R. & Holden, D. W. Functions of the Salmonella pathogenicity island 2 (SPI-2) type III secretion system effectors. Microbiology 158, 1147–1161, doi:10.1099/mic.0.058115-0 (2012).

11 Wang, Y., Huang, H., Sun, M., Zhang, Q. & Guo, D. T3DB: an integrated database for bacterial type III secretion system. BMC Bioinformatics 13, 66, doi:10.1186/1471-2105-13-66 (2012).

12 Arnold, R. et al. Sequence-based prediction of type III secreted proteins. PLoS Pathog 5, e1000376, doi:10.1371/journal.ppat.1000376 (2009).

13 Wang, Y., Zhang, Q., Sun, M. A. & Guo, D. High-accuracy prediction of bacterial type III secreted effectors based on position-specific amino acid composition profiles. Bioinformatics 27, 777–784, doi:10.1093/bioinformatics/btr021 (2011).

14 Dong, X., Lu, X. & Zhang, Z. BEAN 2.0: an integrated web resource for the identification and functional analysis of type III secreted effectors. Database : the journal of biological databases and curation 2015, bav064, doi:10.1093/database/bav064 (2015).

15 Lower, M. & Schneider, G. Prediction of type III secretion signals in genomes of gram-negative bacteria. PloS one 4, e5917, doi:10.1371/journal.pone.0005917 (2009).

16 Ghosh, P. Process of protein transport by the type III secretion system. Microbiology and molecular biology reviews : MMBR 68, 771–795, doi:10.1128/MMBR.68.4.771-795.2004 (2004).

17 McDermott, J. E. et al. Computational prediction of type III and IV secreted effectors in gram-negative bacteria. Infection and immunity 79, 23–32, doi:10.1128/IAI.00537-10 (2011).

18 Altschul, S. F. et al. Gapped BLAST and PSI-BLAST: a new generation of protein database search programs. Nucleic acids research 25, 3389–3402 (1997).

19 Cortes, C. & Vapnik, V. Support-vector networks. Machine learning 20, 273–297 (1995).

20 Gophna, U., Ron, E. Z. & Graur, D. Bacterial type III secretion systems are ancient and evolved by multiple horizontal-transfer events. Gene 312, 151–163 (2003).

21 Medini, D., Covacci, A. & Donati, C. Protein homology network families reveal step-wise diversification of Type III and Type IV secretion systems. PLoS computational biology 2, e173, doi:10.1371/journal.pcbi.0020173 (2006).

22 Radivojac, P. et al. A large-scale evaluation of computational protein function prediction. Nature methods 10, 221–227, doi:10.1038/nmeth.2340 (2013).

23 Goldberg, T. et al. LocTree3 prediction of localization. Nucleic acids research 42, W350–355, doi:10.1093/nar/gku396 (2014).

24 Wang, Y., Sun, M., Bao, H. & White, A. P. T3_MM: A Markov Model Effectively Classifies Bacterial Type III Secretion Signals. PloS one 8, e58173, doi:10.1371/journal.pone.0058173 (2013).

25 Finn, R. D. et al. The Pfam protein families database: towards a more sustainable future. Nucleic acids research 44, D279–285, doi:10.1093/nar/gkv1344 (2016).

26 UniProt Consortum. Reorganizing the protein space at the Universal Protein Resource (UniProt). Nucleic acids research 40, D71–75, doi:10.1093/nar/gkr981 (2012).

27 Zhou, Q., Su, X. & Ning, K. Assessment of quality control approaches for metagenomic data analysis. Scientific reports 4, 6957, doi:10.1038/srep06957 (2014).

28 Konovalova, A., Petters, T. & Sogaard-Andersen, L. Extracellular biology of Myxococcus xanthus. FEMS microbiology reviews 34, 89–106, doi:10.1111/j.1574-6976.2009.00194.x (2010).

29 Pedruzzi, I. et al. HAMAP in 2015: updates to the protein family classification and annotation system. Nucleic acids research 43, D1064–1070, doi:10.1093/nar/gku1002 (2015).

30 Salinero, K. K. et al. Metabolic analysis of the soil microbe Dechloromonas aromatica str. RCB: indications of a surprisingly complex life-style and cryptic anaerobic pathways for aromatic degradation. BMC genomics 10, 351, doi:10.1186/1471-2164-10-351 (2009).

31 Fuerst, J. A., Sambhi, S. K., Paynter, J. L., Hawkins, J. A. & Atherton, J. G. Isolation of a bacterium resembling Pirellula species from primary tissue culture of the giant tiger prawn (Penaeus monodon). Applied and environmental microbiology 57, 3127–3134 (1991).

32 Cayuela, M. L., Elias-Arnanz, M., Penalver-Mellado, M., Padmanabhan, S. & Murillo, F. J. The Stigmatella aurantiaca homolog of Myxococcus xanthus high-mobility-group A-type transcription factor CarD: insights into the functional modules of CarD and their distribution in bacteria. Journal of bacteriology 185, 3527–3537 (2003).

33 Nielsen, P. & Krogh, A. Large-scale prokaryotic gene prediction and comparison to genome annotation. Bioinformatics 21, 4322–4329, doi:10.1093/bioinformatics/bti701 (2005).

34 Okada, N. et al. Identification and characterization of a novel type III secretion system in trh-positive Vibrio parahaemolyticus strain TH3996 reveal genetic lineage and diversity of pathogenic machinery beyond the species level. Infection and immunity 77, 904–913, doi:10.1128/IAI.01184-08 (2009).

35 Lang, J. M., Darling, A. E. & Eisen, J. A. Phylogeny of bacterial and archaeal genomes using conserved genes: supertrees and supermatrices. PloS one 8, e62510, doi:10.1371/journal.pone.0062510 (2013).

36 Macnab, R. M. Type III flagellar protein export and flagellar assembly. Biochimica et biophysica acta 1694, 207–217, doi:10.1016/j.bbamcr.2004.04.005 (2004).

37 Angot, A., Vergunst, A., Genin, S. & Peeters, N. Exploitation of eukaryotic ubiquitin signaling pathways by effectors translocated by bacterial type III and type IV secretion systems. PLoS Pathog 3, e3, doi:10.1371/journal.ppat.0030003 (2007).

38 Chang, J. H. et al. A high-throughput, near-saturating screen for type III effector genes from Pseudomonas syringae. Proc Natl Acad Sci U S A 102, 2549–2554, doi:10.1073/pnas.0409660102 (2005).

39 Greenberg, J. T. & Vinatzer, B. A. Identifying type III effectors of plant pathogens and analyzing their interaction with plant cells. Curr Opin Microbiol 6, 20–28 (2003).

40 Gurlebeck, D., Thieme, F. & Bonas, U. Type III effector proteins from the plant pathogen Xanthomonas and their role in the interaction with the host plant. J Plant Physiol 163, 233–255, doi:10.1016/j.jplph.2005.11.011 (2006).

41 Guttman, D. S. et al. A functional screen for the type III (Hrp) secretome of the plant pathogen Pseudomonas syringae. Science 295, 1722–1726, doi:10.1126/science.295.5560.1722 (2002).

42 Miao, E. A. & Miller, S. I. A conserved amino acid sequence directing intracellular type III secretion by Salmonella typhimurium. Proc Natl Acad Sci U S A 97, 7539–7544 (2000).

43 Sato, H. & Frank, D. W. ExoU is a potent intracellular phospholipase. Mol Microbiol 53, 1279–1290, doi:10.1111/j.1365-2958.2004.04194.x (2004).

44 Tobe, T. et al. An extensive repertoire of type III secretion effectors in Escherichia coli O157 and the role of lambdoid phages in their dissemination. Proc Natl Acad Sci U S A 103, 14941–14946, doi:10.1073/pnas.0604891103 (2006).

45 Mika, S. & Rost, B. UniqueProt: Creating representative protein sequence sets. Nucleic acids research 31, 3789–3791 (2003).

46 Sander, C. & Schneider, R. Database of homology-derived protein structures and the structural meaning of sequence alignment. Proteins 9, 56–68, doi:10.1002/prot.340090107 (1991).

47 Rost, B. Twilight zone of protein sequence alignments. Protein Eng 12, 85–94 (1999).

48 Bairoch, A. & Apweiler, R. The SWISS-PROT protein sequence database and its supplement TrEMBL in 2000. Nucleic acids research 28, 45–48 (2000).

49 Berman, H. M. et al. The Protein Data Bank. Nucleic acids research 28, 235–242 (2000).

50 Frank, E., Hall, M., Trigg, L., Holmes, G. & Witten, I. H. Data mining in bioinformatics using Weka. Bioinformatics 20, 2479–2481, doi:10.1093/bioinformatics/bth261 (2004).

51 Kuang, R. et al. Profile-based string kernels for remote homology detection and motif extraction. Proc IEEE Comput Syst Bioinform Conf, 152–160 (2004).

52 Hamp, T., Goldberg, T. & Rost, B. Accelerating the Original Profile Kernel. PloS one 8, e68459, doi:10.1371/journal.pone.0068459 (2013).

